# COVID-19 patients have increased levels of membrane-associated and soluble CD48

**DOI:** 10.1101/2022.03.18.484843

**Authors:** Hadas Pahima, Ilan Zaffran, Eli Ben-Chetrit, Amir Jarjoui, Pratibha Gaur, Maria Laura Manca, Dana Reichmann, Efrat Orenbuch-Harroch, Ilaria Puxeddu, Carl Zinner, Alexandar Tzankov, Francesca Levi-Schaffer

## Abstract

COVID-19 is a respiratory-centered systemic disorder caused by SARS-CoV-2. The disease can progress into a severe form causing acute lung injury.

CD48 is a co-signaling receptor, existing as both membrane-bound and soluble forms reported to be dysregulated in several inflammatory conditions. Therefore, we reasoned that CD48 could be deregulated in COVID-19 as well.

Here we analyzed CD48 expression in autoptic sections and peripheral blood leukocytes and sera of COVID-19 patients by gene expression profiling (HTG® autoimmune panel), immunohistochemistry, flow cytometry and ELISA.

Lung tissue of COVID-19 patients showed increased CD48 mRNA expression and infiltration of CD48+ lymphocytes. In the peripheral blood, mCD48 was considerably increased on all evaluated cells, and additionally, sCD48 levels were significantly higher in COVID-19 patients independently of disease severity. Considering the alterations of mCD48 and sCD48, a specific role for CD48 in COVID-19 can be assumed, suggesting it as a potential target for therapy.

## Introduction

COVID-19 is a respiratory-centered systemic disorder caused by the severe acute respiratory syndrome (SARS-CoV-2) virus. The first case was described in December 2019 and by the beginning of February 2022, 380,321,615 cases and 5,680,741 deaths have been reported worldwide according to the World Health Organization (WHO). Most SARS-CoV-2 infected individuals will experience mild symptoms or even be asymptomatic(Wiersinga *et al*, 2020). However, the disease can progress into a severe form causing acute lung injury (ALI), mainly diffuse alveolar damage (DAD) with thromboinflammation, immunopathology, and cytokine storm syndrome(Bösmüller *et al*, 2021). Interleukin-6 (IL-6) is one of the cytokines involved in progressive and severe disease and was suggested together with C-reactive protein (CRP) to be a useful predictor for mechanical ventilation(Herold *et al*, 2020).

CD48, (SLAMF2), is a glycosylphosphatidylinositol (GpI) activating or co-activating receptor, belonging to the SLAM family. It is expressed on most hematopoietic cells as is its high affinity ligand 2B4 (CD244). As several other GpIs, CD48 exists both as a membrane bound (mCD48) and a soluble (sCD48) form(Pahima *et al*, 2019). In moderate asthma, mCD48 is significantly increased on peripheral blood eosinophils and B cells, while in severe asthma in addition to B cells mCD48 is significantly increased on T-cells, NK-cells, and monocytes. sCD48 levels are significantly increased in the sera of mild asthmatic patients, compared to healthy donors while severe asthma patientstreated with glucocorticosteroids display significantly decreased levels of sCD48(Gangwar *et al*, 2017). However, we concluded that the decrease in sCD48 was due to asthma severity rather than to a direct effect of the drug(Gangwar & Levi-Schaffer, 2016). Importantly, CD48 increase in asthma was neither correlated with Th2 inflammation biomarkers such as IL-33, IL-5, IgE, and eosinophils numbers, nor with tobacco smoking or BMI(Breuer *et al*, 2018). Interestingly, sCD48 levels are found to be augmented in the sera of individuals infected with the Epstein-Bar Virus(Smith *et al*, 1997) and varicella, and measles infections are accompanied by increased expression of mCD48 on monocytes and lymphocytes(Katsuura *et al*, 1998). Additionally, the owl monkey cytomegalovirus (CMV) was reported to utilize the decoy property of sCD48 and to produce sCD48 homologues to escape the immune response(Pérez-Carmona *et al*, 2015).

Based on all these evidences, we reasoned that CD48 expression could be differentially dysregulated in COVID-19. In the present work we report on data obtained by *in situ* analysis of CD48 expression in lung specimens collected *post-mortem* and by analysis of mCD48 expression on peripheral blood cells and of sCD48 in serum of COVID-19 patients with various disease severities.

## Results

### Laboratory findings

Routine blood tests of COVID-19 patients and healthy controls are shown in Fig 1 and Table 1. As expected, COVID-19 patients displayed peripheral blood neutrophilia, lymphocytopenia and eosinopenia and significant upregulation of CRP and IL-6 levels (Fig 1A-C) that were associated with disease severity (Fig 1F-G). As in other reports critical and/or severe patients showed a significant decrease in hemoglobin levels (Fig 1H), and increased levels of D-dimer and ferritin levels, albeit non-significant (Fig S2).

**Fig 1:**
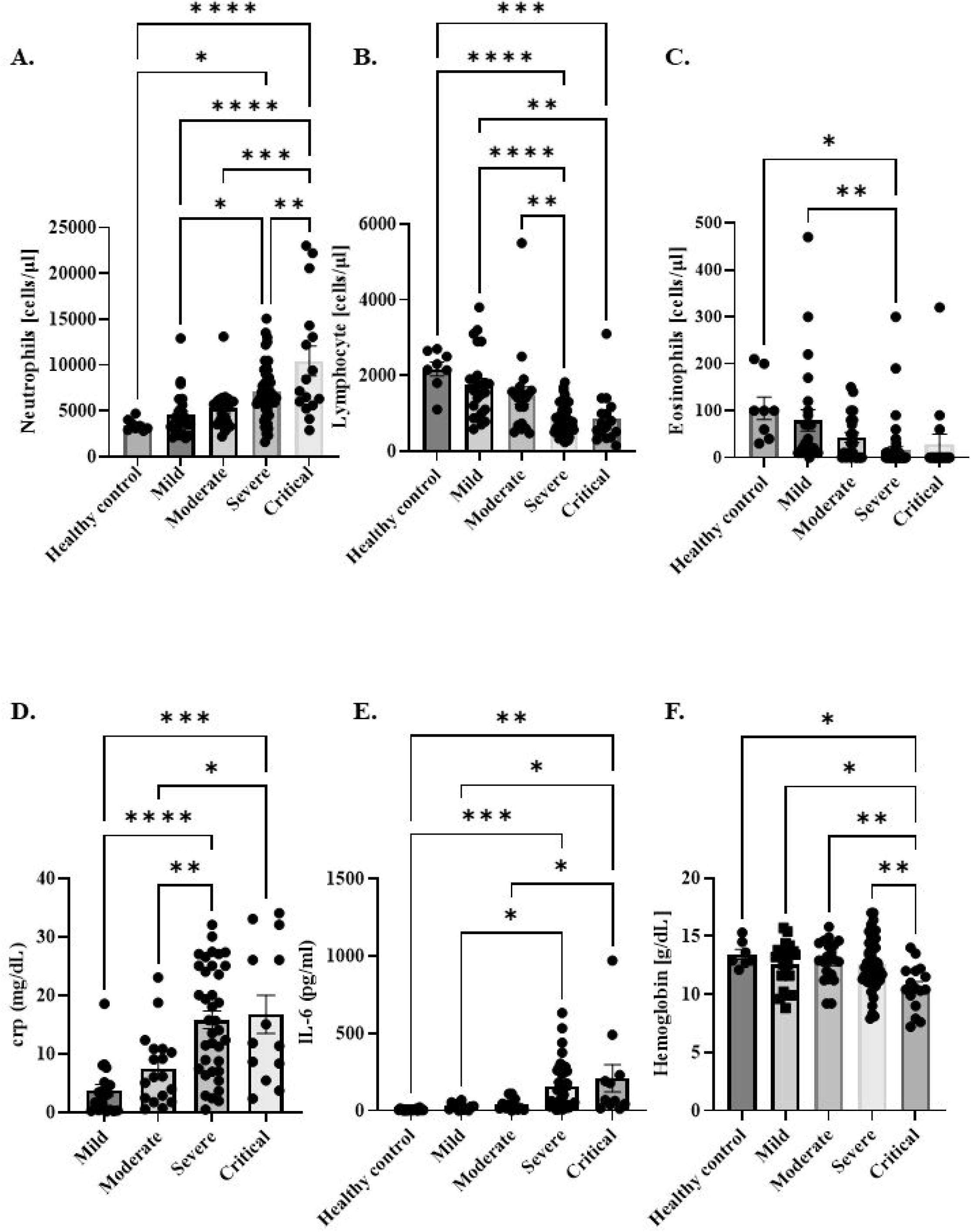
Laboratory findings in COVID-19 patients and in healthy controls: **(A)** Neutrophils (cells/μl); **(B)** Lymphocytes (cells/μl); **(C)** Eosinophils (cells/μl); **(D)** CRP levels (mg/dL); **(E)** IL-6 (pg/ml) levels; **(F)** Hemoglobin (g/dL) from COVID-19 patients and/or healthy controls. Data are shown as the mean ± SEM. *P < 0.05, **P < 0.01, ***P < 0.001, ****P < 0.0001

**Table 1:** Laboratory findings of active patients of COVID-19, with different disease severities. Data presented as mean ± SEM.

|  | Ferritin |  | D-dimer |  | Creatinine |  | CRP |  |
| --- | --- | --- | --- | --- | --- | --- | --- | --- |
|  | mean | n | mean | n | mean | n | mean | n |
| Mild | 533.5 ± 151.2 | 17 | 742.3 ± 128.0 | 19 | 1.263 ± 0.29 | 24 | 4.051 ± 0.89 | 25 |
| Moderate | 544.8 ± 307.7 | 6 | 944.7 ± 350.6 | 11 | 0.787 ± 0.067 | 16 | 6.749 ± 1.47 | 16 |
| Severe | 749.2 ± 123.4 | 32 | 4026 ± 1259 | 38 | 5.910 ± 2.411 | 49 | 13.69 ± 1.34 | 49 |
| Critical | 3220 ± 1868 | 13 | 6822 ± 3210 | 14 | 6.974 ± 5.240 | 16 | 15.71 ± 2.80 | 16 |

### COVID-19 lung tissues display significantly higher CD48 mRNA levels and increased infiltration of CD48+ lymphocytes as compared to other lung pathologies

To assess the expression levels of CD48 in COVID-19 lung tissues, a transcriptomic analysis was performed. The mRNA levels of *CD48* were found to be significantly up-regulated in COVID-19 lungs in comparison to non-COVID-19 DAD, and arterial hypertension controls (Fig 2A).

**Fig 2:**
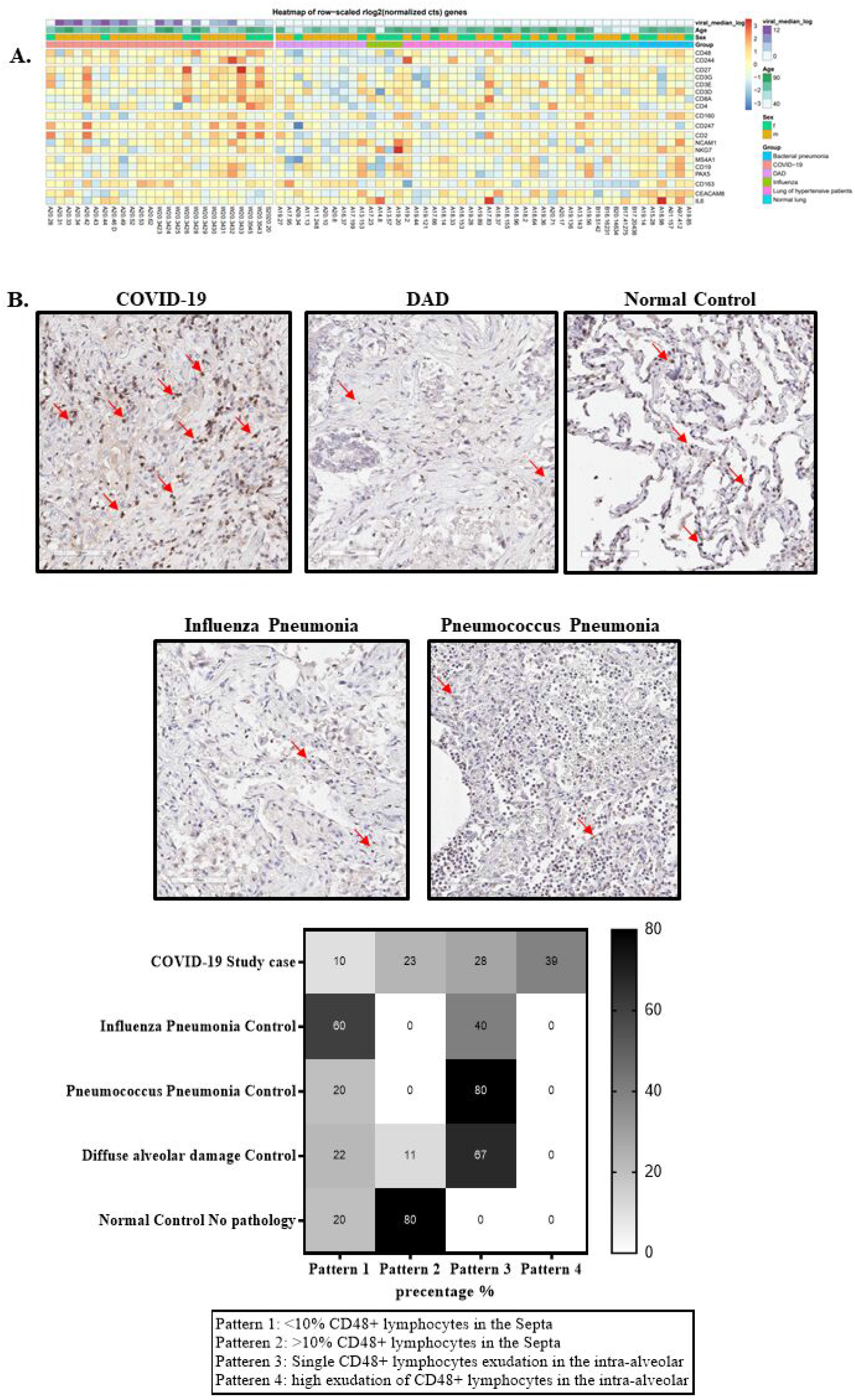
GEP and IHC staining of lung specimens of post-mortem COVID-19 patients, other pathologies, and healthy tissue: **(A)** GEP showed gene expression differences between COVID-19 samples and other pathologies as presented by the heatmap; **(B)** representative images of CD48 tissue staining of either COVID-19, DAD, healthy tissue, influenza pneumonia or pneumococcus pneumonia, X40; **(C)** A semi-quantitative analysis of the different patterns.

When mRNA levels of *CD48* were compared with immune cell-specific genes’ expression an association was found between CD48 and expression genes related to T-cells and NK-cells, but not to B-cells (Fig 2A). mRNA levels of 2B4 were evaluated as well and found not to be significantly different in COVID-19 in comparison to the other diseases (Fig 2A).

Consistently, IHC CD48 staining patterns showed a general increase of “pattern four” in COVID-19 in comparison to the other lung pathologies (Fig 2B). Importantly, CD48 positive lymphocytes were the predominantly infiltrating cells in COVID-19 lung tissues (Fig 2B). Of note, non-COVID-19 DAD lungs showed mostly “pattern three”, even though they displayed significant amounts of CD48 negative intra-alveolar immune cell infiltration.

### Increased CD48 on leukocytes and in serum of COVID-19 patients

Next, we evaluated mCD48 expression on peripheral blood T-cells, B-cells, NK-cells, monocytes, and neutrophils from COVID-19 patients with different disease severity and from healthy controls (FC). Interestingly, the expression levels of mCD48 were ubiquitously higher on patient cells in comparison to healthy controls’ cells (Fig 3A-E, Table 2). The highest increase was found on monocytes (2.2 folds) and NK-cells (2 folds). CD48 expression on neutrophils was almost undetectable and increased only by 1.3-fold. (Fig 3E, Table 2). Interestingly, mCD48 expression enhancement was not linked to disease severity, except for B-cells, where mCD48 expression was significantly increased in mild-moderate patients in comparison to severe-critical patients’ cells (S3A).

**Fig 3:**
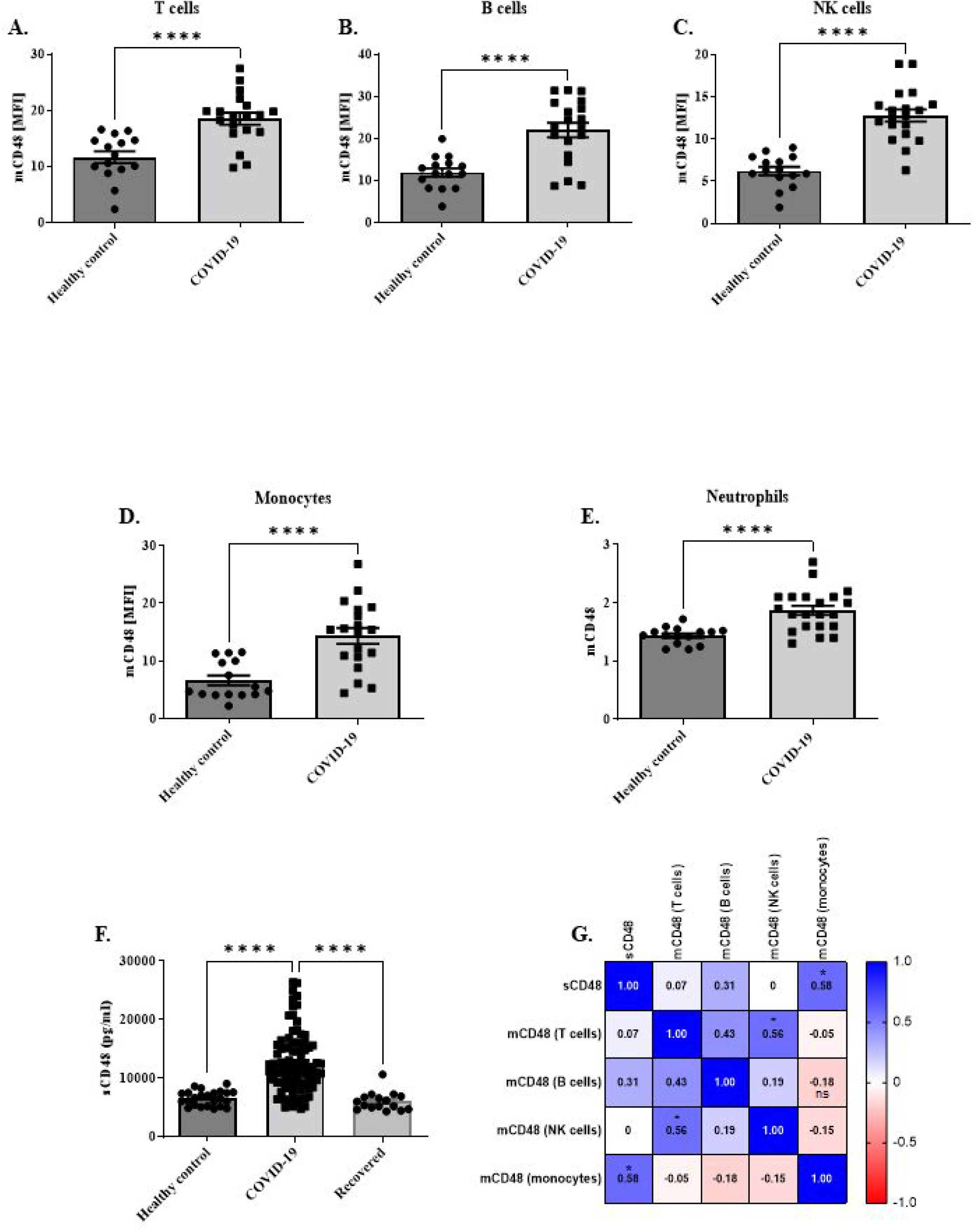
Membrane associated and soluble and CD48 levels of COVID-19 patients and healthy controls: mCD48 expression on **(A)** T cells; **(B)** B cells; **(C)** NK cells; **(D)** Monocytes; **(E)** Neutrophils from COVID-19 patients and HC identified by staining with their specific cell surface markers; **(F)** sCD48 levels in the sera of COVID-19 patients, healthy controls and COVID-19 recovered individuals; **(G)** Multiplex correlation assay. Data are shown as the mean ± SEM. *P < 0.05, **P < 0.01, ***P < 0.001, ****P < 0.0001

**Table 2:** mCD48 expression levels (MFI) on peripheral blood leukocytes of aged-matched COVID-19 active patients and healthy controls. Data presented as mean ± SEM

|  | Healthy controls | COVID-19 |
| --- | --- | --- |
| Age (mean) | 54 ± 4 | 57 ± 4 |
| mCD48 on NK cells (mean) | 6 ± 0.5 | 13 ± 1 |
| mCD48 on T cells (mean) | 12 ± 1 | 18 ± 1 |
| mCD48 on B cells (mean) | 12 ± 1 | 22 ± 2 |
| mCD48 on Monocytes (mean) | 7 ± 1 | 14 ± 1 |
| mCD48 on Neutrophils (mean) | 1.44 ± 0 | 1.88 ± 0 |
| n | 15 | 19 |

Moreover, the sCD48 levels in the sera of 88 COVID-19 patients with different disease severity were significantly augmented in comparison to healthy controls (24). sCD48 levels were 2-fold higher in COVID-19 patients (12543 ± 541 pg/ml) compared to the healthy control group (6555 ± 258 pg/ml) (Fig 3F, Table 3) (p-value < 0.0001). However, as seen in Fig S2A, this increment was not associated with disease severity since similar values were obtained for mild (11859 ± 1303 pg/ml), moderate (9690 ± 857 pg/ml), severe (13882 ± 751 pg/ml), and critical (12056 ± 1580 pg/ml) patients. It is noteworthy that sCD48 levels returned to the ones of healthy controls once patients recovered from the disease (5961 ± 376) (Fig 3F, Table 3).

**Table 3:** sCD48 levels in the sera of aged-matched active or recovered COVID-19 patients and healthy controls. Data presented as mean ± SEM

|  | Healthy controls | COVID-19 all | Mild | Moderate | Severe | Critical | Recovered |
| --- | --- | --- | --- | --- | --- | --- | --- |
| sCD48 pg/ml (mean) | 6555 ± 258 | 12511 ± 546 | 11859 ± 1303 | 9690 ± 8567 | 13882 ± 751 | 12056 ± 1580 | 5997 ± 398 |
| Age (mean) | 62 ± 3 | 63 ± 2 | 56 ± 6 | 54 ± 4 | 67 ± 2 | 69 ± 2 | 49 ± 3 |
| n | 24 | 88 | 18 | 14 | 42 | 14 | 16 |

A multiplex correlation analysis found a moderate positive correlation (r: 0.58; p-value: 0.026) between sCD48 levels and mCD48 expression on monocytes of COVID-19 patients (Fig 3G). It is noteworthy that a similar correlation was not found for sCD48 and mCD48 on monocytes of healthy controls (S2C). Correlation analysis also detected a positive correlation (r: 0.56; p-value: 0.031) between mCD48 levels on T cells and mCD48 levels on NK-cells and (Fig 2G).

Next, we wondered whether there is a correlation between sCD48 levels and inflammatory markers known to be increased in parallel with COVID-19 severity. A moderate positive correlation (r=0.42, p-value: 0.0005) was found between IL-6 levels and sCD48 levels in the sera of COVID-19 patients (Fig S2G). Finally, a positive weak correlation was found between sCD48 and CRP levels (r=0.24; p-value: 0.0260) (Fig S2H).

### m2B4 levels are upregulated on NK-cells and monocytes in COVID-19 patients

As mentioned above 2B4 is the high-affinity ligand of CD48. Therefore, 2B4 levels were evaluated on blood leukocytes from 8 COVID-19 patients (3 mild, 1 moderate and 4 severe) and 12 healthy controls. 2B4 levels were found to be slightly, but significantly elevated on NK-cells and monocytes (4.854 ± 0.48; 7.775 ± 0.8, respectively) in comparison to healthy controls (3.72 ± 0.19; 3.85 ± 0.5, respectively), while on B-cells, T-cells, and neutrophils 2B4 levels were undetectable (data not shown) (Fig 4A-B). Surprisingly, high levels of m2B4 on NK-cells were positively correlated with high levels of mCD48 on either T-cells (r: 0.8573; p-value: 0.0065), NK-cells (r: 0.9036; p-value: 0.0072) or monocytes (r: 0.7289; p-value: 0.047) (Fig 4C-E). Noteworthy, the correlation between m2B4 and mCD48 on NK-cells was found also in healthy controls (r: 0.8629; p-value: 0.0005), while the other correlations were detected only in the patients’ cells (Fig 4F-H).

**Fig 4:**
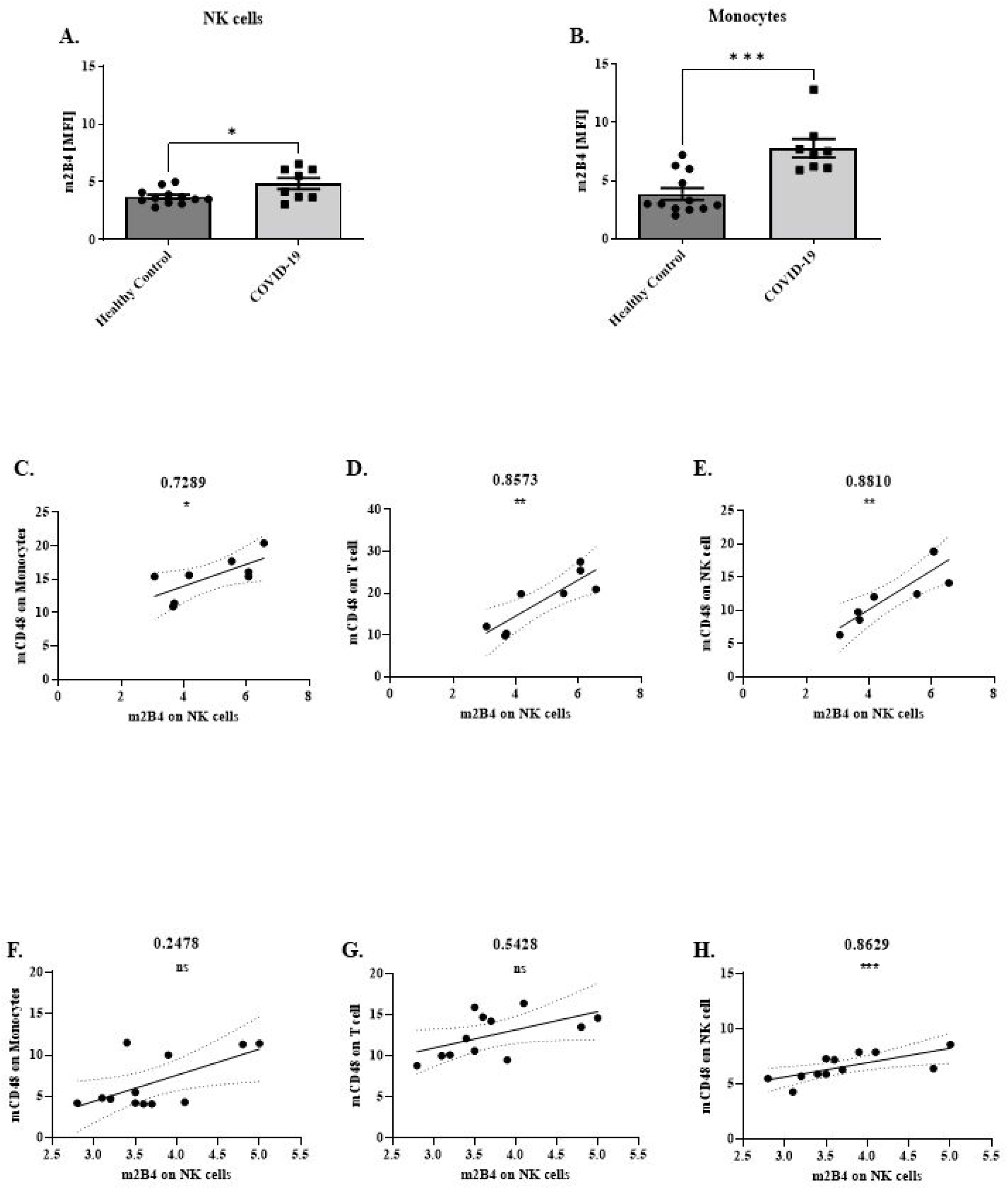
Membrane associated 2B4 levels of COVID-19 patients and healthy controls: m2B4 expression on **(A)** NK cells; **(B)** Monocytes from COVID-19 patients and HC identified by staining with their specific cell surface markers. Data are shown as the mean ± SEM. *P < 0.05, **P < 0.01, ***P < 0.001, ****P < 0.0001. Spearman correlation analysis between m2B4 on NK cells and mCD48 on **(C)** Monocytes; **(D)** T cells; and **(E)** NK cells of COVID-19 patients or **(F)** Monocytes; **(G)** T cells; and **(H)** NK cells of HC.

## Discussion

The present study provides a comprehensive examination of the effect of SARS-CoV-2 on the expression of CD48 in the lungs and in the peripheral blood of COVID-19 patients. To investigate the influence of SARS-CoV-2 in lung CD48 expression we initially assessed lung tissue obtained *post-mortem* from COVID-19 patients and from other lung pathologies such as non-COVID-19 DAD, influenza pneumonia, pneumococcal pneumonia, and from healthy individuals.

The ALI caused by COVID-19 was reported to have many morphological similarities to other non-COVID-19 DAD. However, some COVID-19 peculiarities are observed such as massive capillarostasis, intussusceptive angiogenesis and a massive increased NETosis(Bösmüller *et al*, 2021). Here we report on an additional parameter, rather specific for COVID-19 DAD, i.e. the significant upregulation of CD48 mRNA in the affected lungs, compared to non-COVID-19 DAD. Importantly, the lung tissues were obtained from deceased COVID-19 patients during the first wave of the pandemic, who did not receive dexamethasone for their disease. This finding suggests that CD48 levels may increase specifically due to either SARS-CoV-2 infection or still consequently because of lung injury. Notably, the augmented mRNA levels of *CD48* correlated with increased mRNA levels of CD8+ T-cell-characteristic genes and - to a lesser extent - of NK-cell genes (Fig 1A). Of note, *CD48* mRNA increment was not associated with high or low SARS-CoV-2 viral burden (data not shown).

Interestingly, Huang et al showed that *in-vitro* infection of pluripotent stem cell-derived human lung alveolar type 2 cells with SARS-CoV-2 causes a decrease in the mRNA levels of *CD48*(Huang *et al*, 2020). These observations could bolster our data of increased mRNA levels in infiltrating immune cells rather than on structural cells.

Unlike *CD48*, mRNA levels of its high-affinity ligand, *2B4*, did not show significant changes when compared to the other lung pathologies studied and healthy lung tissues. An exception were the lungs of influenza virus infected patients that showed a decrease of *2B4* mRNA. The expression of *CD48* mRNA in the bronchoalveolar lavage fluid of COVID-19 was reported by Desterke et al(Desterke *et al*, 2020). In their study, CD48 enhancement was related to disease severity and was associated with a CD14-CD16 subpopulation. In our study, we didn’t find a particular correlation between *CD48* mRNA and monocyte and/or macrophages markers.in the lungs

Concordant with the GEP, we found that also at the protein level, CD48-positive immune cell levels were significantly increased in COVID-19. Moreover, COVID-19 affected lungs were the only ones characterized by CD48+ intra-alveolar lymphocytosis. This was in contrast to the fact that both COVID-19 and non-COVID-19 DAD contained similar amounts of intra-alveolar immune cells. Even though no differences were reported in lymphocyte infiltration in the lungs of COVID-19 and influenza-infected individuals in general(Ackermann *et al*, 2020), currently no CD48+ lymphocytes were found in the intra-alveolar space of influenza-infected patients. All these imply a specific importance of CD48 in COVID-19.

Consistently, in further analyses, SARS-CoV-2 infected individuals showed a significant increase of mCD48 on most of the peripheral blood leukocytes in comparison to healthy controls, regardless of disease severity. To the best of our knowledge, this is the first time that mCD48 levels have been found enhanced on peripheral blood leukocytes following a respiratory viral infection. We previously found that mCD48 levels are altered in asthma(Gangwar *et al*, 2017) and COPD [unpublished data]. mCD48 levels in COVID-19 patients might be elevated due to the concomitant increased levels of INFγ(Akbari *et al*, 2020), known to upregulate CD48 expression, or due to other still undiscovered factors. Notably, virus infections seem to influence expression levels of mCD48 as on B-cells obtained from mononucleosis patients mCD48 is elevated(Thorley-Lawson *et al*, 1982), while on peripheral blood CD4+ T cells from T-lymphotropic virus 1 (HTLV-1) infected patients it is decreased(Ezinne *et al*, 2014), and on NK-cells of herpes simplex virus (HSV) infected patients it remains stable(Lenart *et al*, 2021). Furthermore, mCD48 expression is reported to be elevated in CD4+ T-cells infected *in vitro* with the human immunodeficiency virus (HIV)(Tremblay-McLean *et al*, 2017).

Similarly, to mCD48, sCD48 was found to be significantly higher in the sera of COVID-19 patients, with a positive correlation between sCD48 and mCD48 levels on monocytes, suggesting monocytes as primary cells releasing sCD48. Interestingly, this increase was independent of disease severity and returned to baseline upon recovery. Noteworthy, the effect of treatment on sCD48 was evaluated in mild-moderate patients, since only in this group untreated patients were available.

However, no significant difference was found between treated and untreated patients (data not shown) suggesting that, as in asthma, also in COVID-19 OCS do not affect CD48. Furthermore, since no differences in sCD48 levels were found between blood samples obtained from the three different waves (data not shown), we may assume that sCD48 release is not affected by the different SARS-CoV-2 variants and treatment strategy variations.

In this study we measured as accepted markers of COVID-19 severity, IL-6 and CRP. A moderate or weak positive correlation was found between IL-6 or CRP, respectively, and sCD48 levels.

We have previously reported that sCD48 is a decoy receptor for both 2B4 and *Staphylococcus aureus* enterotoxin B (SEB) on human peripheral blood eosinophils and in SEB-induced peritonitis in mice(Gangwar & Levi-Schaffer, 2016). We can therefore hypothesize that sCD48 in COVID-19 might downregulate the immune response against SARS-CoV-2 by masking its ligand 2B4. Interestingly, CMV was reported to produce a sCD48 homolog, named A43, which inhibits NK-cell function through binding to 2B4, allowing CMV to escape the immune response(Martínez-Vicente *et al*, 2019). 2B4 expression levels, on both NK-cells and monocytes are significantly higher than healthy controls, as also detected following *in-vitro* infection with influenza virus and *in-vivo* administration of influenza vaccine(Jost *et al*, 2011). Remarkably, in our study higher levels of 2B4 on NK-cells were correlated with higher levels of CD48 on T-cells and monocytes (Fig 3C-D). The engagement between CD48 on monocytes and 2B4 on NK-cells might inhibit the functionality of the latter cells, as described in hepatocellular carcinoma(Wu *et al*, 2013). At least in our collective, this correlation was specific to COVID-19 patients and was not found in healthy controls. The strong positive correlation between m2B4 and mCD48 on NK-cells is most likely related to homeostasis, as it was found on both COVID-19 and healthy controls.

In summary, we infer that the increase of mCD48 on the lung-infiltrating lymphocytes, which is paralleled on the peripheral blood lymphocytes and monocytes, along with the increased levels of sCD48 in the serum of COVID-19 patients might indicate a decrease of anti-viral immune response. Moreover, since sCD48 levels are increased already in patients with mild symptoms and remain high even in critical patients, CD48 can be deemed a marker of symptomatic SARS-CoV-2 infection.

## Methods

### Study Cohort

Lung tissues from 28 patients who died of COVID-19 (Appendix Table S1; Appendix Table S2) during the first disease wave in Switzerland (March-May 2020) were organized into a tissue microarray (TMA) format, as reported previously(Tzankov *et al*, 2021; Menter *et al*, 2020). The original paraffin blocks of each patient were punched three times with a 1mm core needle. For the purposes of the study, areas of superposed pneumonia were excluded, in order to obtain informative samples from the areas of primary COVID-19-related damage. The results were compared to the results obtained from similarly arrayed cases of 5 influenza viral pneumonias, 5 pneumococcal pneumonias, 5 other causes (non-infectious and non-COVID-19) diffuse alveolar damage (DAD; referred to as “other cause DAD”), and 5 normal lungs. Tissue collection was approved by the Ethics committee of Northern and Central Switzerland (study ID 2020-00969). None COVID-19 patient, except for one treated with low dose prednisone, has been treated with OCS; especially none has received dexamethasone as per treatment protocols established later in the pandemics.

Blood was obtained from a total of 111 COVID-19 active (Appendix Table S3, Appendix Table S4) and 18 recovered (Appendix Table S5) patients, hospitalized either in Shaare Zedek Medical Center (86 patients; P1-P86) or Hadassah University Hospital (5 patients; P87-P91 and 2 recovered; R1-R2) (Jerusalem, Israel) and in Pisa University Hospital (20 patients; P92-P11 and 15 recovered; R3-R18) (Pisa, Italy) and from 26 (Appendix Table S6), age and sex matched, healthy controls in Hadassah University Hospital (Jerusalem, Israel). Patients were classified for disease severity according to WHO guidelines, and the study included 26 patients with mild, 20 patients with moderate, 49 patients with severe, and 16 with critical disease, respectively. Subjects’ peripheral blood collection and experiments were approved by the Helsinki committee of each hospital.

### Gene Expression Programming (GEP) of lung tissue

GEP of the lung tissue obtained at autopsy and used for the TMA construction was performed by HTG according to established protocols (https://www.htgmolecular.com/assets/htg/resources/BR-05-HTG-EdgeSeq-System.pdf). Lysates from samples were run on the HTG EdgeSeq Processor (HTG Molecular Diagnostics, Tucson, AZ, USA) using the HTG EdgeSeq Immune Response Panel with an excess of nuclease protection probes (NPPs) complimentary to their target. S1 nuclease then removed un-hybridized probes and RNAs leaving behind NPPs hybridized to their targets in a 1-to-1 ratio. Samples were individually barcoded using a 16-cycle PCR reaction to add adapters and molecular barcodes, individually purified using AMPure XP beads (Beckman Coulter, Brea, CA, USA) and quantitated using a KAPA Library Quantification kit (KAPA Biosystems, Wilmington, MA, USA). Libraries were sequenced on the Illumina SEQUENCER platform (Illumina, San Diego, CA, USA) for quantification. Quality control, standardization and normalization were performed by HTG and provided to the investigators. Quality control criteria as determined by the manufacturer (percentage of overall reads allocated to the positive process control probe per sample <28%, read depth ≥750000, relative standard deviation of reads of each probe within a sample >0.094) were met for all samples.

Data were first analyzed by the HTG online tool (https://reveal.htgmolecular.com/), including manual analysis of the genes of interest. A PCA analysis was then performed using the pcomp function in R©, version 4.0.3 (R-Project for Statistical Computing, Vienna, Austria) and differential expression analysis of COVID-19 cases against controls was conducted with the DESeq2 package using default settings.

Count estimates were normalized with the median ratio method, and low-quality samples were excluded from analysis. Prior to the heatmap visualization, the normalized counts were further log2-transformed using a robust variance stabilization. The heatmap was produced with the pheatmap package. The column clusters of the samples as well as the row clusters of the significant genes were obtained by hierarchical clustering with complete linkage and a Euclidean distance metric. The Wald test-statistic was utilized, and p-values were adjusted for false discoveries. Adjusted p-values <0.05 and | log2 (fold change) | > 1 were considered significant and included in the analysis.

### Immunohistochemistry

With the exception of CD48 (see below) immunohistochemistry (IHC) was performed on the TMAs using the automated staining system Benchmark XT (Roche/Ventana Medical Systems, Tucson, USA) as per ISO15189 accredited standard operating procedure of the Institute of Pathology at the University Hospital Basel. *In-situ* hybridization for SARS-CoV-2 was performed as previously reported(Reinhold *et al*, 2021). For CD48 IHC, slides were stained by Dako autostainer after deparaffinization by warming up to 75°C. Antigen retrieval was achieved at 95°C with the Ultra Cell Conditioner for 8 min. Slides were then incubated for 40 min at 37°C with 36 μg/mL anti-CD48 antibody (EPR4108; ab134049; Abcam, Cambridge, UK), washed, and counterstained with hematoxylin (Gill II).

Stained sections were scanned using the Aperio AT2 scaner (Leica, Wetzlar, Germany). Images were visualized via Aperio ScanScope Console and analyzed manually according to the following parameters: less than 10% CD48 positive lymphocytes in the alveolar septa were categorized as “pattern one”, while more the 10% cells were considered as “pattern two”. “Pattern three” was when subtle CD48 positive lymphocyte exudation in the intra-alveolar space was noticed, and “pattern four” corresponded to heavy intra-alveolar exudation of CD48 positive cells.

### Leukocyte isolation from peripheral blood of COVID-19 patients and controls

The white blood cell fraction from peripheral blood of COVID-19 patients and healthy controls was isolated according to a previously described protocol(Gangwar *et al*, 2017). The obtained granulocytes and mononuclear cells were washed and resuspended in the same medium as granulocytes. For serum collection, venous blood (1-2 ml) was withdrawn in non-heparinized tubes and centrifuged (2000 rpm, 10 min, 4°C). The collected serum was stored at −80°C.

### Human peripheral blood leukocytes double staining

Isolated fractions of granulocytes and mononuclear cells (2×10^5^/100 μl of 0.1% BSA/PBS) were incubated in 96 U-bottom plates (Thermo Fisher Scientific Inc., Waltham, MA) on ice. For flow cytometry (FC) double staining, cells were blocked (5% goat serum, 0.1% BSA/PBS 15 min on ice) and incubated with either FITC-anti-human CD3 (clone HIT3a; BioLegend, San Diego, CA) and PE-anti-human CD56/NCAM (HCD56; BioLegend, San Diego, CA) for T cells and NK cells, FITC-anti-human CD20 (2H7; BioLegend, San Diego, CA) for B cells, FITC anti-human CD14 (HCD14; BioLegend, San Diego, CA) for monocytes, or FITC anti-human CD16 (3G8; Santa Cruz Biotechnology, Dallas, TX) for neutrophils. For the analysis of cell surface molecules cells were stained with APC anti-human CD48 (MEM-102; Abcam, Cambridge, UK); APC-anti human 2B4/CD244 (2-69; BioLegend, San Diego, CA); or with the isotype control APC mouse IgG1 (MOPC-21; BioLegend, San Diego, CA). Thereafter, cells were washed twice (250g, 5 min, 4°C) with 0.1% BSA/PBS, and FC data were acquired by BD LSR II Flow Cytometer and analyzed with FlowJo software (Tree Star, OR, USA). Cells populations were gated according to physical parameters and surface marker specific staining (Fig S1). MFIs were determined by dividing the fluorescence intensity of the specific mAb by that of the isotype control. Fig S1b shows a representative histogram of CD48 staining.

### Measurement of soluble CD48 and IL-6 levels in serum

Serum levels of soluble CD48 (sCD48) and IL-6 were quantified using commercially available ELISA kits (human CD48 ELISA kit; DEIA237 Sensitivity: 9.38pg/ml; Creative Diagnostic, Shirley, NY, USA; and Deluxe set Human IL-6, cat: 430504 Sensitivity: 4pg/ml; BioLegend, San Diego, CA, USA) according to the manufacturer’s instructions. IL-6 and CD48 ELISAs were performed in duplicates using undiluted serum and 1:40 diluted, respectively, and values were calculated based on a recombinant standard curve.

### Statistical analysis

Continuous data are expressed as mean ± standard error of the mean (SEM), or median and interquartile range. Variables with a skewed distribution (by the Shapiro-Wilk test) were log-transformed for use in parametric testing. Statistical comparisons between experimental groups were performed using one-way analysis of variance (ANOVA) and post-hoc Tukey’s multiple comparison tests. For fewer than three experimental groups, Student’s unpaired two-tailed t-test was employed.

Associations were evaluated by Spearman correlation testing. Data were analyzed by Prism version 9.0 (GraphPad Software, San Diego, California, USA). A p-value less than 0.05 was considered statistically significant for all analyses.

## ACKNOWLEDGEMENTS

This work was supported by grants from the Israel Science Foundation (ISF, 3933/19) to Francesca Levi-Schaffer. Francesca Levi-Schaffer is affiliated with the Adolph and Klara Brettler Center for Molecular Pharmacology and Therapeutics at the School of Pharmacy of The Hebrew University of Jerusalem. The authors thank Dr. Hadas Segev-Yekutiel (The Core Research Facility, The Faculty of Medicine, The Hebrew University of Jerusalem) who provided advice on the FC analysis and technical help and to all the donors who donated the blood for the study.

## AUTHOR CONTRIBUTIONS

H.P planned and performed experiments, analyzed the data, prepared the figures, and wrote the manuscript. I.Z planned and performed experiments, analyzed the data, prepared the figures, and edited the manuscript. P.G assisted with the experiments. A.J provided blood, biochemical and clinical information from IL patients and assisted in the interpretation of the results. M.L.M assisted with the statistical analysis. D.R assistance with biochemical assays. E.O.H provided blood, biochemical and clinical information from IL patients. I.P provided sera, biochemical and clinical information from IT patients and assisted in the interpretation of the results. C.Z performed the analysis of the GEP assay. E.B.C Organized the blood collection and provided blood samples, biochemical and clinical information from IL patients and assisted in the interpretation of the results. A.T Provided the autoptic lung sections for both RNA analysis and tissue array IHC, critically read and edited the manuscript. F.L.S designed and supervised the study, analyzed the data, advised, reviewed, and edited the manuscript.

## Supplementary figures and legends

**Fig EV1:** Gating and identification of leucocytes from peripheral blood by flow cytometry. **(A)** Peripheral blood leucocytes were first separated by their physical parameters (FSC and SSC) and granulocytes, lymphocytes, and mononuclear cells were identified and gated. From the mononuclear cells gate monocytes (CD14+) were identified, from the lymphocytes gate, B cells (CD20+), T cells (CD56-/CD3+), and NK cells (CD56+/CD3−) were identified, and from the granulocyte gate neutrophils (CD16+) were identified. **(B)** representative histogram of CD48 and Isotype control intensity on each cell type.

**Fig EV2:** Laboratory findings of COVID-19 patients and healthy controls: **(A)** Ferritin; (ng/ml) (**B)** d-Dimer; (ng/ml) **(C)** cre (mg/dL) levels; **(D)** Monocytes (cells/μl); **(E)** Basophils (cells/μl); **(F)** WBC (cells/μl); **(G)** platelets (cells/μl) from COVID-19 patients and/or HC. Data are shown as the mean ± SEM. *P < 0.05, **P < 0.01, ***P < 0.001, ****P < 0.0001.

**Fig EV3**: Membrane associated and soluble and CD48 levels of COVID-19 patients and healthy controls: **(A)** mCD48 expression on peripheral blood leukocytes of COVID-19 patients with different disease severity; **(B)** sCD48 levels in the sera. Data are shown as the mean ± SEM. *P < 0.05, **P < 0.01, ***P < 0.001, ****P < 0.0001

